# A novel vimentin-stabilizing urea compound MXC-017 ameliorates DSS-induced colitis and radiation enteropathy in mice

**DOI:** 10.64898/2026.08.30.748104

**Authors:** Ling He, Linda Azizi, Carlos Calderon, Sophia Jin, Talia Parker, Rikhil Seth, Xiaohong Chen, Hui Ding, Michael Jung, Frank Pajonk

## Abstract

Ulcerative colitis (UC) and radiation enteropathy involve intestinal epithelial injury, barrier dysfunction, and inflammation, but effective treatments remain limited. This study evaluated MXC-017, a novel vimentin-targeting urea compound, in mouse models of dextran sulfate sodium (DSS)-induced colitis and radiation-induced enteropathy.

Acute colitis was induced in C57BL/6 mice using 3.5% DSS for seven days, followed by regular water for seven days. Radiation enteropathy was induced by 13 Gy total abdominal irradiation. Mice received MXC-017 (150 mg/kg) or vehicle. Disease activity, intestinal permeability, inflammatory and epithelial markers, and histopathology were assessed. MXC-017’s effects on cancer stem cell frequency, sphere formation, and migration were also examined in PC-3 and DU-145 prostate cancer cells.

MXC-017 reduced DSS-induced colitis severity, accelerated weight recovery, lowered disease activity, partially preserved colon length, and restored barrier function. It also reduced proinflammatory cytokines, macrophage infiltration, epithelial injury, and goblet cell loss while preserving epithelial proliferation and markers of intestinal stem cell function and tight-junction integrity. Following irradiation, MXC-017 improved weight recovery, reduced intestinal permeability, preserved epithelial architecture, and partially mitigated villus shortening.

Importantly, MXC-017 did not protect prostate cancer stem cells from radiation. Instead, it reduced stem cell frequency, sphere-forming capacity, and cancer cell migration. These findings support vimentin targeting with MXC-017 as a potential treatment for UC and radiation-induced intestinal toxicity and as an adjunct to radiotherapy for pelvic and abdominal malignancies.

## Introduction

In 2019 inflammatory bowel disease (IBD) affected 4.9 million patients worldwide with the USA and China reporting the most cases. The incidence has steadily risen over the past three decades making IBD a major health burden and driver of health care costs [1]. IBD can be classified into two main types: Crohn’s disease and ulcerative colitis (UC). Crohn’s disease can affect any part of the gastrointestinal tract and is characterized by transmural inflammation, while UC is limited to the colon and rectum and is characterized by mucosal inflammation. The exact cause of IBD is complex and includes genetic disposition, environmental, immune, and microbiological factors [2].

Current treatment options for UC remain limited and primarily rely on anti-inflammatory or immunosuppressive strategies, including 5-aminosalicylates (5-ASAs), corticosteroids, biologics targeting TNFα or integrins, Janus Kinase (JAK) inhibitors, interleukin inhibitors, and other immunomodulatory agents [3–7]. Severe, therapy-refractory cases can require surgical intervention.

Histopathological, UC is characterized by disruption of the intestinal epithelial barrier and depletion of the protective mucus layer, resulting in increased intestinal permeability and sustained inflammatory responses that contribute to disease progression [8–12].

Likewise, exposure of the intestines to ionizing radiation, either in the context of accidental exposure or during radiotherapy of abdominal malignancies, can cause a similar pathology, summarized as radiation enteropathy [13]. The most widely accepted mechanism is the elimination of intestinal stem cells with subsequent loss of the epithelial layer, transmigration of microbes, leading to local infections or even sepsis [14–17]. There are very few treatment options to mitigate the effects of accidental radiation exposure or -aside from exposure avoidance-to protect the intestines against radiation effects during radiotherapy [18, 19]. So far, no victim of accidental radiation exposures who developed an acute gastro-intestinal radiation syndrome has survived [20–22] and localized radiation exposure of the intestines as a part of cancer therapy can lead to chronic proctitis, strictures and fistulas [23–25].

In this study we used the Dextran Sulfate Sodium (DSS)-induced mouse model of colitis [26, 27] and a total abdominal irradiation model to test if a novel vimentin-targeting urea compound could mitigate colitis symptoms and radiation enteropathy.

## Material and Methods

### Animals

Male and female 7-9-week-old C57BL/6 mice were re-derived, bred and maintained in a pathogen-free environment in the American Association of Laboratory Animal Care-accredited Animal Facilities of the Division of Laboratory Animal Medicine, University of California, Los Angeles. The protocols were approved by the University of California Los Angeles’ Institutional Animal Care and Use Committee (IACUC).

### Cell lines

PC3 and DU-145 cell lines were grown in log-phase in DMEM supplemented with 10% fetal bovine serum (Thermo Fisher Scientific, MA), 1 % penicillin, and streptomycin. All cells were grown in a humidified atmosphere at 37°C with 5% CO_2_. The identity of both cell lines was confirmed by DNA fingerprinting (Laragen, Culver City, CA). Cell lines were routinely tested for mycoplasma infection (#G238, Applied Biological Materials, Ferndale, WA).

### DSS-Induced Acute Colitis and Drug Treatment

Acute colitis was induced by administering 3.5% dextran sulfate sodium (DSS, molecular weight 36-50 kDa, Sigma) in the drinking water for 7 consecutive days, followed by regular drinking water for an additional 7 days. MXC-017 (N-[(4-piperidin-1-ylsulfonylphenyl)methyl]indole-1-carboxamide) was synthesized in-house and freshly prepared each day by dissolving in 1% DMSO with corn oil and administered intraperitoneally from day 1 to day 14 (150 mg/kg). Mice in the control group received solvent control (1% DMSO in corn oil). Body weight and Disease Activity Index (DAI) scores were recorded daily throughout the 14-day period. The DAI score was calculated as the sum of individual scores (ranging from 0 to 4) for body weight loss, stool consistency, and fecal bleeding **(Supplementary Table 1)**. In compliance with ethical guidelines, mice were euthanized if body weight loss exceeded 20%.

### Irradiation

Cells were irradiated at RT using an experimental X-ray irradiator (Gulmay Medical Inc. Atlanta, GA) at a dose rate of 5.3985 Gy/min. Control samples were sham-irradiated.

For total abdominal irradiation animals were anesthetized prior to irradiation with an intra-peritoneal injection of 30 µL of a ketamine (100 mg/mL, Phoenix, MO) and xylazine (20 mg/mL, AnaSed, IL) mixture (4:1). Mice were irradiated in groups of 3 using an experimental X-ray irradiator (Gulmay Medical Inc. Atlanta, GA) at a dose rate of 6.704 Gy/min for the time required to apply a prescribed dose of 13 Gy with the whole body except for the abdomen shielded. The X-ray beam was operated at 300 kV and hardened using a 4 mm Be, a 3 mm Al, and a 1.5 mm Cu filter. NIST-traceable dosimetry was performed using a micro-ionization chamber. Mice were euthanized on day 6 after irradiation.

### Serum FITC-Dextran Intestinal Permeability Assay

Intestinal permeability was assessed using a fluorescein isothiocyanate-dextran (FITC– dextran; 4 kDa, Sigma) assay. Mice were fasted for 4 hours with free access to water, followed by oral gavage with FITC-dextran (400 mg/kg body weight) freshly prepared in sterile PBS (80 mg/mL). After 4 hours, blood samples were collected by cardiac puncture and allowed to clot at room temperature for 30 minutes. Serum was obtained by centrifugation at 2,000 *× g* for 10 minutes and stored at −80 °C until analysis. Serum fluorescence was measured using a microplate reader (SpectraMax iD3, Molecular Devices, San Jose, CA, USA; excitation 485 nm, emission 530 nm). FITC-dextran concentrations were calculated from a standard curve prepared by spiking known amounts of FITC-dextran (0-320 µg/mL) into pooled naive mouse serum. Increased serum FITC-dextran levels were interpreted as indicative of compromised intestinal barrier integrity.

### Quantitative Reverse Transcription-PCR

Total RNA was isolated from distal mouse colon tissue and small intestine tissue using TRIZOL Reagent (Invitrogen, Waltham, MA). cDNA synthesis was carried out using the SuperScript Reverse Transcription IV (Invitrogen). Quantitative PCR was performed in the QuantStudio^TM^ 3 Real-Time PCR System (Applied Biosystems, Carlsbad, CA, USA) using the PowerUp^TM^ SYBRTM Green Master Mix (Applied Biosystems). Ct for each gene was determined after normalization to GAPDH and ^ΔΔ^Ct was calculated relative to the designated reference sample. Gene expression values were then set equal to 2^−ΔΔCt^ as described by the manufacturer of the kit (Applied Biosystems). All PCR primers were synthesized by Invitrogen. GAPDH was used as a housekeeping gene (for primer sequences see **Supplementary Table 2**).

### Histopathological Analysis

Freshly collected colon and small intestine tissues were collected separately, washed and arranged in a Swiss-roll configuration, fixed in formalin, dehydrated, and embedded in paraffin. Sections were cut at 4-µm thickness and stained with hematoxylin and eosin (H&E), and alcian blue for histopathological evaluation.

For immunohistochemistry analysis, the sections were deparaffinized in xylene and rehydrated through a graded ethanol series. The slides were incubated in 0.1% Triton-X in PBS for 10 minutes. Antigen retrieval was performed using Heat Induced Epitope Retrieval in a citrate buffer (10 mM sodium citrate, 0.05% tween20, pH 6) with heating to 95°C in a steamer for 25 minutes. After cooling down, the slides were blocked with 10% goat serum plus 1% BSA at room temperature for 30 minutes and then incubated with the primary antibody against Ki67 (1:400; #12202s; Cell Signaling Technology; Danvers, MA), E-cadherin (1:200; #14472s; Cell Signaling Technology), Occludin (1:200; #91131s; Cell Signaling Technology), ZO-1 (1:200; ab221547, Abcam), F4/80 (1:200; #70076t; Cell Signaling Technology), Vimentin (1:200; #5741s; Cell Signaling Technology), TNFα (1:200; #11948s; Cell Signaling Technology), or IL-1β (1:100; #12507s; Cell Signaling Technology) overnight at 4°C. The next day, the slides were rinsed with PBS and then incubated with ready-to-use IHC detection reagent (Cell Signaling Technology) at room temperature for 1 h, rinsed, and then incubated with DAB (Cell Signaling Technology) for 1-5 minutes. Tissues were counterstained with Harris modified Hematoxylin (Fisher scientific, Waltham, MA) for 30 seconds, dehydrated via an alcohol gradient (ethanol 25%, 50%, 70%, 90%, 100%) and soaked twice into Xylene. A drop of Premount mounting media (Fisher Scientific) was added on the top of each section before covering up with a coverslip.

### Alcian Blue staining

Colon sections were deparaffinized, hydrated and acid-treated in 3% glacial acetic acid for 3 minutes. Alcian Blue Solution (pH 2.5) for was applied 30 minutes at room temperature. Slides were then rinsed in distilled water for 2 minutes to remove excess dye, and nuclei were counterstained with Nuclear Fast Red solution for 5 minutes. After rinsing in running tap water for 1 minute, slides were dehydrated through graded alcohols, cleared in xylene, and mounted.

### Immunofluorescence staining

PC-3 or DU-145 cells were trypsinized and plated onto 2-well chamber slides (Lab-Tek, Thermo Fisher Scientific) at a density of 10,000 cells/ well for 24 hours prior to staining. Cells were then fixed with 4% paraformaldehyde for 10 minutes at room temperature, permeabilized with 0.1% Triton X-100 for 10 minutes, and blocked with 10% goat serum for 30 minutes at room temperature. Cells were then incubated with vimentin antibody (1:200; #5741s; Cell Signaling Technology) overnight at 4°C. The next day, cells were washed with PBS and incubated with Alexa Fluor 488 goat anti-rabbit IgG (1:1000, A-11008, Thermo Fisher Scientific) for 1 hour at room temperature in the dark. Cells were then washed with Pbs and counterstained with Hoechst 33342 (1:5000, Thermo Fisher Scientific) for 10 minutes at room temperature in the dark. Finally, cells were washed with PBS and mounted on slides with Prolong Gold Antifade Reagent (Thermo Fisher Scientific). Images were acquired using a confocal microscope (Zeiss).

### Sphere Formation Assay (SFA) and Extreme Limiting Dilution Analysis (ELDA)

PC-3 or DU-145 cells were trypsinized and serially diluted in non-tissue culture-treated 96-well plates under serum-free conditions at a range from 1 to 512 cells/well. Growth factors (EGF and bFGF) were supplemented every two days. Prostate spheres were counted 7 days later and presented as a percentage of the initial number of cells plated. Prostate cancer stem cell frequencies were calculated using the ELDA software included in the statmod (vs 1.5.2) package in R [28].

### Immunofluorescent staining

Cells were plated onto the 2-chamber cell culture slides (#154461, ThermoFisher). For vimentin staining, cells were fixed, permeabilized, blocked and stained with monoclonal vimentin antibody (#5741s, cell signaling technology, 1:100) followed by Alexa Fluor 488 Goat Anti-rabbit IgG (H/L) secondary antibody (1:1,000 (Invitrogen)) and nuclear counterstaining. Fluorescent images were acquired using a confocal microscope (Zeiss).

### Migration Assay

PC3 or DU-145 monolayers were plated on 3 cm Petri dishes in 10% FBS medium. 24 hours later, the cells were serum starved overnight in medium containing 1% FBS. The next day, cells were trypsinized and plated onto cell culture inserts with an 8 µm pore size (Corning Inc., NY) at 100K cells/well. To facilitate cell migration in the chambers, an FBS gradient was created by filling the bottom chambers with medium containing 10% FBS supplemented with either 10 µM MXC-017 or an equivalent volume of DMSO as the vehicle control. After 20-24 hours of incubation, cells were fixed using 10% formalin and washed with PBS. Using a cotton swab, non-migrated cells on the upper side of the membrane were gently removed and migrated/attached cells at the bottom of the membrane were stained with 1% crystal violet. Subsequently, images were taken with a digital microscope (BZ-9000, Keyence, Itasca, IL), and migrated cells were counted using the Image J software.

Statistics

All data shown are represented as mean ± standard error mean (SEM) of at least 3 biologically independent experiments. A *p*-value of ≤0.05 in one-way ANOVA followed by Tukey’s multiple comparison test in Graphpad Prism (vs 11.1.0) indicated a statistically significant difference.

## Results

### MXC-017 attenuates DSS-induced colitis symptoms

Exposure to DSS is a well-established model of colitis [26, 27]. In this study, we used the DSS-induced mouse model of colitis and total abdominal irradiation to test if a novel urea compound could mitigate colitis symptoms and radiation enteropathy. When mice were exposed to DSS at 3.5% for 7 days in their drinking water **(Figure 1A)**, they developed colitis symptoms, including weight loss, diarrhea and rectal bleeding. However, when DSS exposure was combined with MXC-017 (**Figure 1B**) treatment, symptoms were milder. Weight loss and diarrhea were similar initially, but after day 7, mice treated with MXC-017 started to gain weight **(Figure 2A)** and had less diarrhea and rectal bleeding than the DSS-only group. Consequently, the DAI score was lower starting on day 6 and returned to normal by day 14 in the DSS/MXC-017 group while the DSS-only group continued to have high DAI scores **(Figure 2B)**. Colon lengths were reduced in the DSS-only group compared to controls and partially restored in the DSS/MXC-017 group (5.75 cm vs 5 cm, *p*-value = 0.035; **Figure 2C/D**).

**Figure 1.**
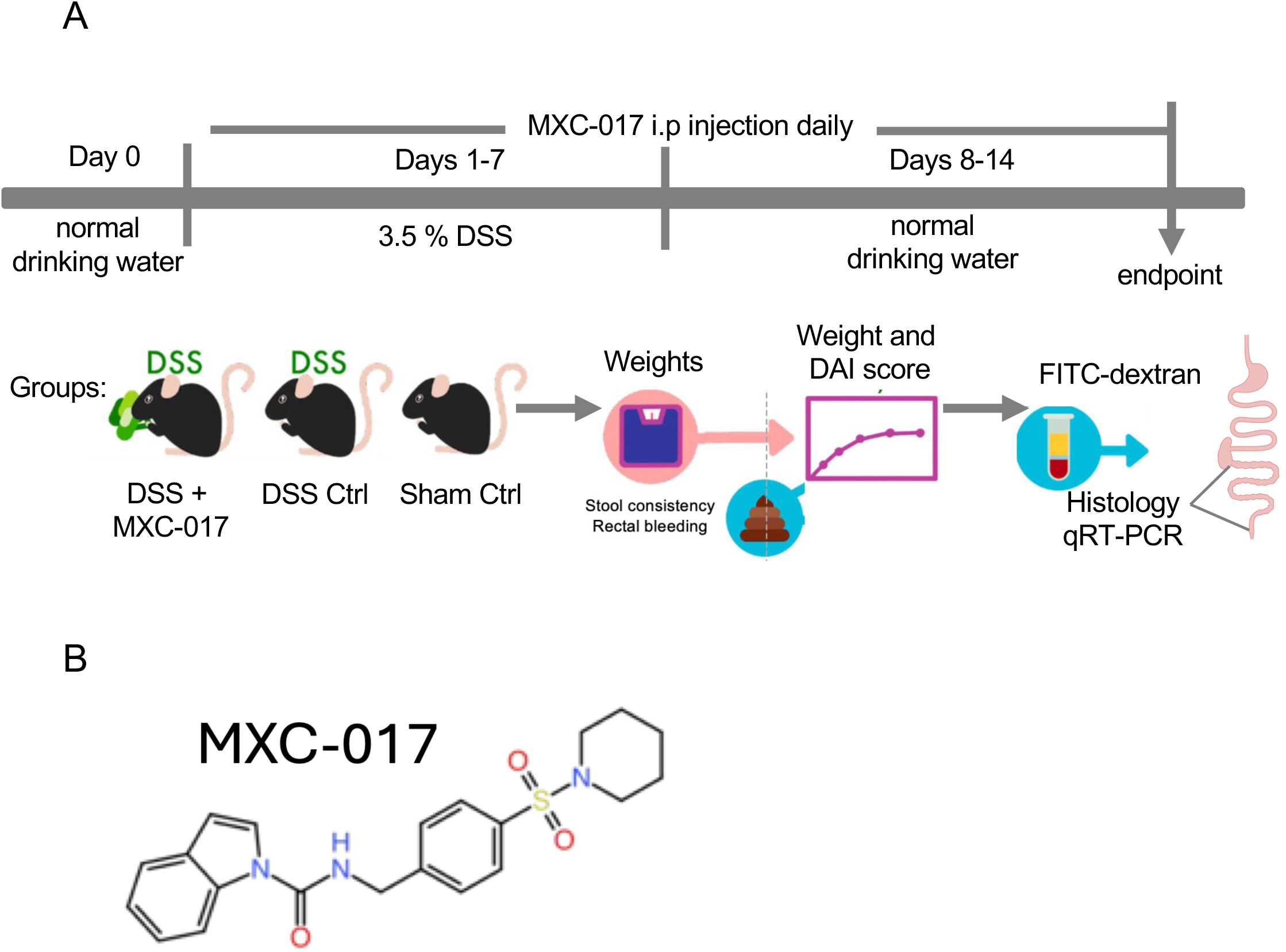
**(A)** Experimental design and therapeutic effects of MXC-017 in a DSS-induced colitis model. (**B**) Chemical structure of MXC-017.

**Figure 2.**
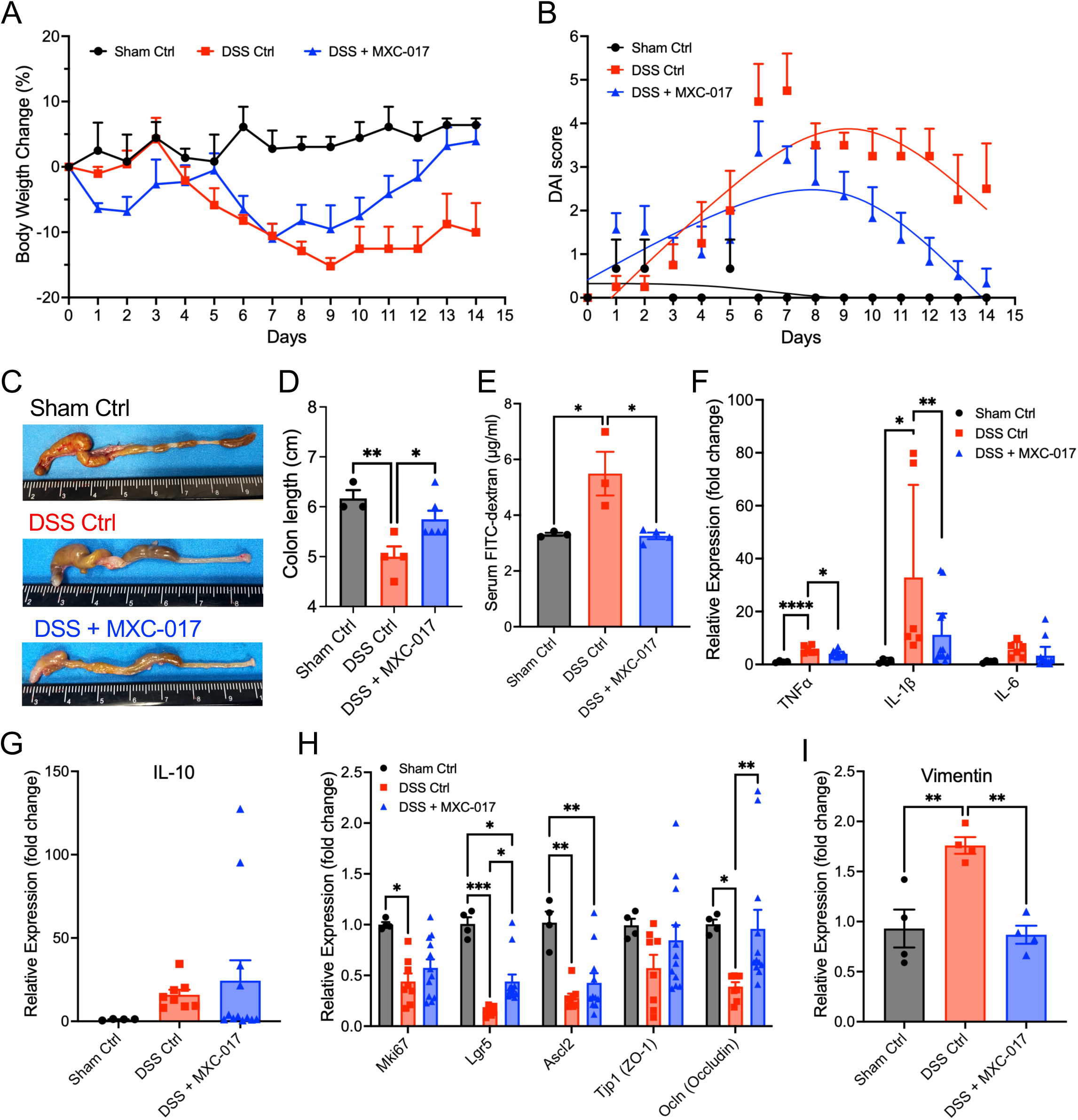
**(A)** Changes in body weight over time in sham control, DSS-treated, and DSS + MXC-017-treated mice. **(B)** Disease Activity Index (DAI) scores in the indicated treatment groups. **(C)** Representative images of colons harvested from Sham control, DSS-treated, and DSS+MXC-017-treated mice. **(D)** Quantification of colon length at the experimental endpoint. **(E)** Serum FITC-dextran levels as a measure of intestinal permeability and barrier function. **(F/G)** Relative mRNA expression of the pro-inflammatory cytokines IL-1β, TNF-α, and IL-6 and the anti-inflammatory cytokine IL-10 in colon tissue collected at the experimental endpoint. **(H)** Relative mRNA expression of the proliferation marker Ki-67, intestinal stem cell markers Lgr5 and Ascl2, ang tight junction proteins ZO-1 and Occludin**. (I)** Relative Vimentin expression in colon tissue at the experimental endpoint. Data were obtained from sham control, DSS-treated, and DSS+MXC-017-treated mice. 3-6 animals were included in each group. P-values were calculated using one-way ANOVA. * p-value < 0.05, ** p-value < 0.01, *** p-value < 0.001, **** p-value < 0.0001.

Next, we tested if MXC-017 could mitigate the loss of the intestinal barrier function measuring FITC-labeled dextran serum levels after oral administration of FITC-dextran 14 days after the start of the DSS treatment. FITC-dextran serum levels were significantly higher in the DSS-only group compared to controls, thus indicating a loss of the intestinal barrier function (5.49 vs 3.31 µg/ml, *p*-value = 0.0213). In contrast, FITC-dextran serum levels were significantly lower in the DSS/MXC-017 group compared to the DSS-only group (3.26 vs 5.49 µg/ml, *p*-value = 0.0137) and similar to those of the control group **(Figure 2E)**. DSS treatment led to a significant increase in proinflammatory TNFα and IL-1β mRNA expression levels in colon tissue compared to controls and treatment with MXC-017 significantly reduced DSS-induced TNFα and IL-1β mRNA expression. A similar trend was observed for IL-6 expression but did not reach statistical significance **(Figure 2F)**. In contrast, mRNA level of the anti-inflammatory cytokine IL-10 was increased in both the DSS control and DSS+MXC-017 groups relative to sham control, but these changes did not reach statistical significance **(Figure 2G)**.

To further assess epithelial integrity and regenerative capacity, we examined the expression of proliferation-, stem cell-, and barrier-associated genes. DSS treatment significantly reduced the expression of Ki67, Lgr5, Ascl2, and the tight junction marker Occludin, consistent with impaired epithelial proliferation, intestinal stem cell function, and barrier integrity. Treatment with MXC-017 partially restored the expression of these markers toward control levels **(Figure 2H)**. As we previously identified vimentin as the molecular target of MXC-017 [29], we also evaluated vimentin expression in colon tissue. DSS exposure significantly increased vimentin expression, indicative of tissue injury and stromal activation, whereas MXC-017 attenuated this response **(Figure 2I)**, suggesting that modulation of vimentin-associated pathways may contribute to its protective effects in colitis.

Consistent with the development of systemic inflammation, DSS treatment resulted in significant splenomegaly compared with control animals **(Supplementary Figure 1A/B)**. However, MXC-017 treatment did not significantly reduce DSS-induced spleen enlargement. To determine whether MXC-017 altered regulatory immune responses within the colon, we further examined the expression of Foxp3 and Tgfb1, two markers associated with regulatory T-cell activity and immune suppression. Neither marker showed significant differences among the experimental groups **(Supplementary Figure 1C)**, suggesting that the protective effects of MXC-017 are unlikely to be mediated through major alterations in Treg-associated pathways in the colon.

H&E staining of colon tissue revealed an increase in inflammatory cell infiltration in the DSS control and DSS+MXC-017 groups compared with Sham controls. DSS treatment caused substantial disruption of the epithelial architecture, including crypt damage and mucosal erosion, whereas these pathological changes were attenuated by MXC-017 treatment **(Figure 3A)**. Consistent with these findings, alcian blue staining demonstrated a marked depletion of goblet cells and reduced mucin content following DSS treatment, indicating the impairment of mucus barrier integrity. Treatment with MXC-017 partially preserved alcian blue-positive goblet cells and mucin staining, suggesting improved maintenance of the intestinal mucus layer and epithelial barrier function **(Figure 3B)**.

**Figure 3.**
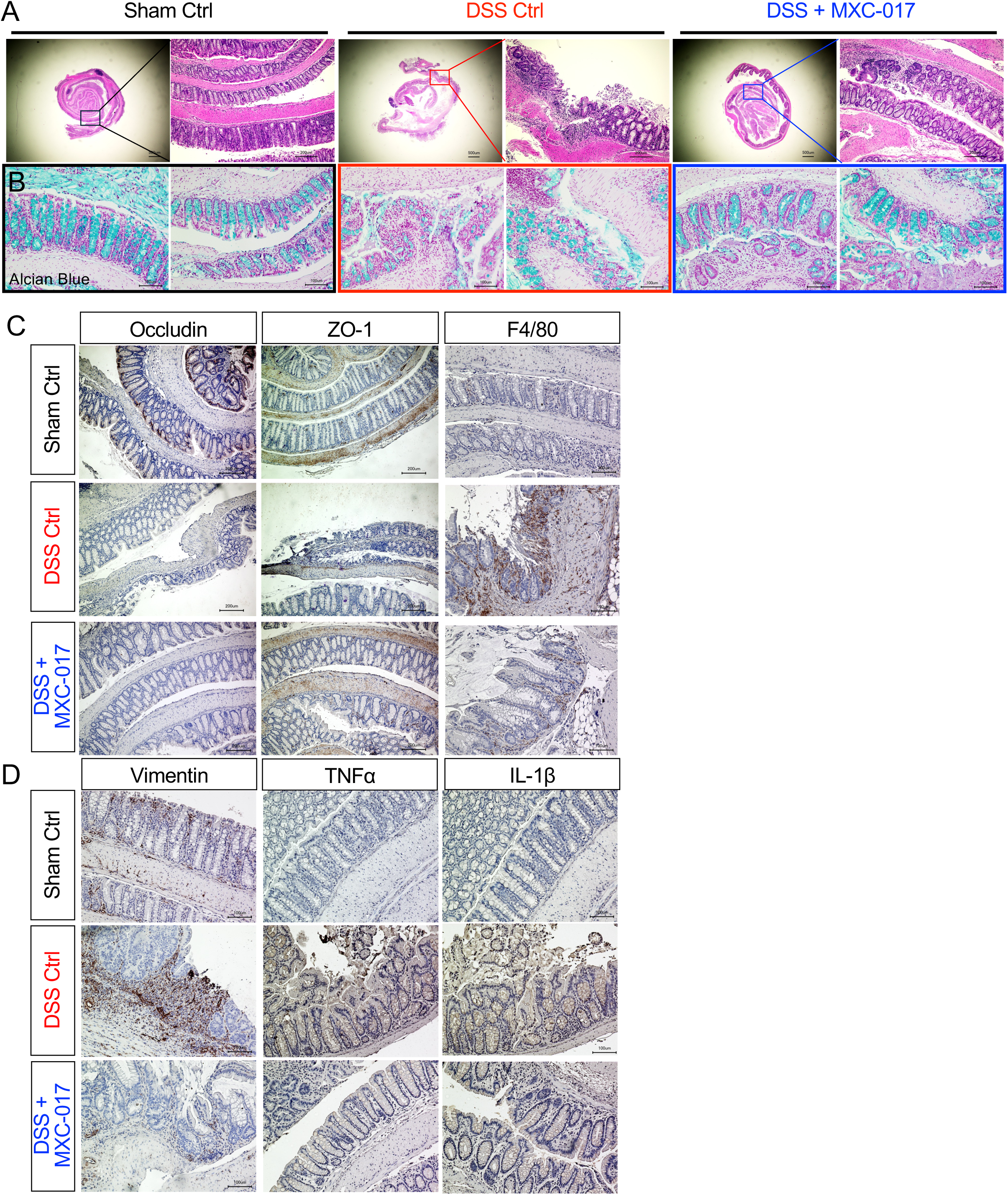
**(A)** Representative H&E-stained Swiss-roll colon sections obtained at the experimental endpoint. Inserts show higher-magnification views of the indicated region of interest (ROI). **(B)** Representative Alcian blue staining of colon tissues. **(C/D)** Representative immunohistochemical staining of Occludin, ZO-1, TNF-α, IL-1β, vimentin, and F4/80 in colon tissues.

The number of Occludin-positive was diminished in the DSS-treated animals compared with Sham controls and not affected by MXC-017 treatment. In contrast, zonula occludens-1 (ZO-1) staining demonstrated a marked depletion of ZO-1-positive cells in the DSS-treated animals compared with Sham controls. However, MXC-017 treatment preserved the number of ZO-1-positive cells in the DSS-treated group, indicating that MXC-017 treatment preserves tight junctions of epithelial barrier (**Figure 3C**).

Following DSS treatment, colon tissue was infiltrated with F4/80-positive macrophages, consistent with a pro-inflammatory response. This infiltration was reduced in the DSS+MXC-017 group and reached almost the same level as the Sham controls (**Figure 3C**). This pro-inflammatory response to DSS was further confirmed by the increased expression of TNFα and IL-1β, both produced by macrophages [30], and this was again reversed by the addition of MXC-017 (**Figure 3D**). Lastly, we stained against the intermediate filament vimentin, which we previously identified as the molecular target of MXC-017 [29]. DSS treatment led to increased expression of vimentin in the colonic epithelium and the addition of MXC-017 reversed this effect (**Figure 3D**), matching the mRNA expression data (**Figure 2I**).

### MXC-017 mitigates radiation-induced enteropathy

To explore if MXC-017 would also mitigate radiation-induced enteropathy in the colon and small intestines, mice received total abdominal irradiation with a single dose of 13 Gy. This dose of radiation is non-lethal but sufficient to induce severe radiation enteropathy in mice [31]. A schematic overview of the experimental design is shown in **Figure 4A**.

**Figure 4.**
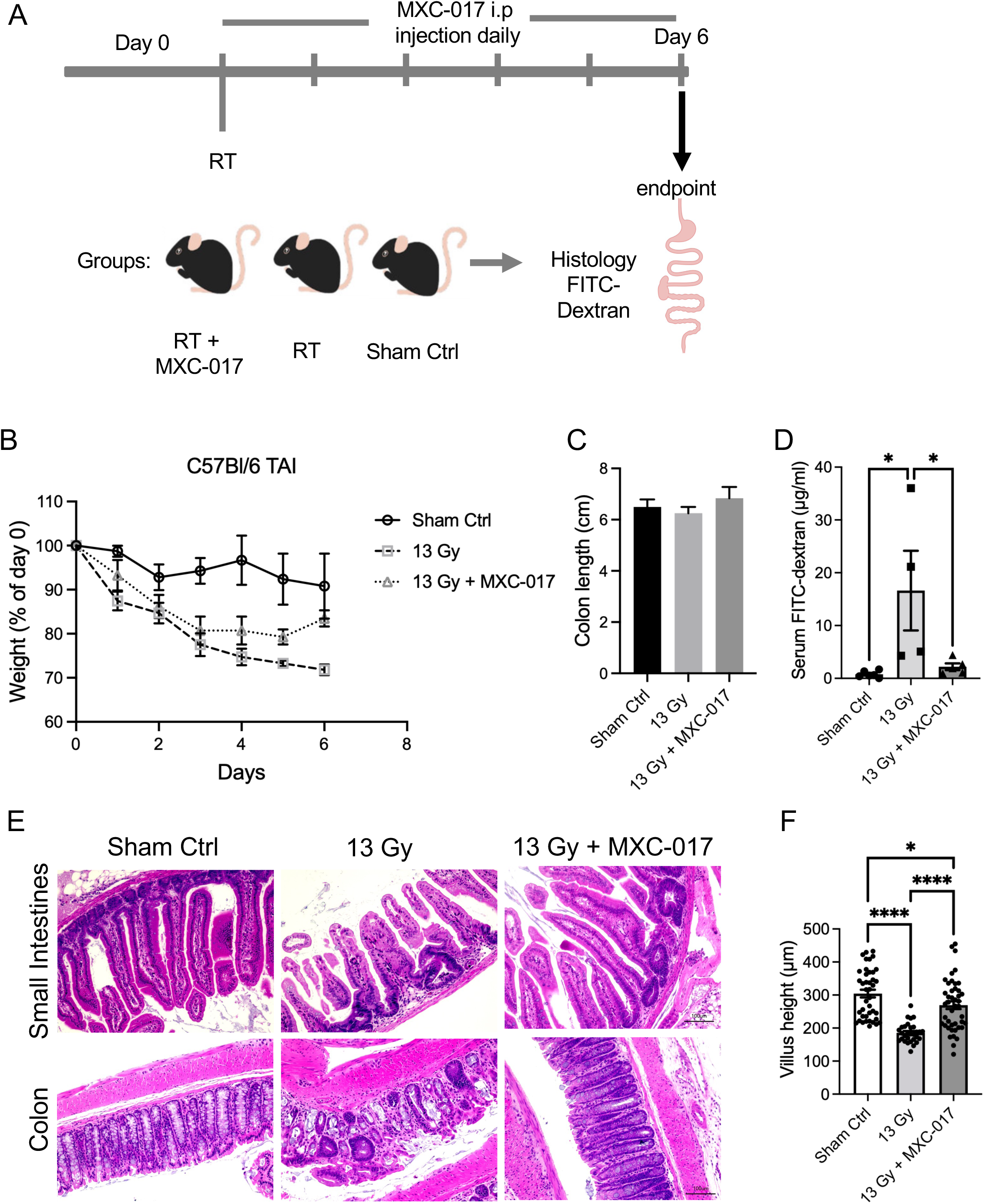
(**A**) Experimental design. (**B**) weight curve of mice treated with total abdominal radiation (13 Gy) or total abdominal radiation plus MXC-017 compared to sham irradiated animals. (**C**) Colon lengths of mice treated with total abdominal radiation (13cGy) or total abdominal radiation plus MXC-017 compared to sham irradiated animals. (**D**) Serum FITC-dextran levels of mice treated with total abdominal radiation or total abdominal radiation plus MXC-017 compared to sham irradiated animals. (**E**) Representative H&E staining of small intestine and colon tissues. Abdominal radiation disrupted the intestinal barrier integrity, and this was ameliorated by MXC-017. (**F**) Radiation-induced shortening of small intestinal villi was ameliorated by MXC-017. 4-6 animals were included in each group. P-values were calculated using one-way ANOVA. * p-value < 0.05, **** p-value < 0.0001.

Radiation exposure resulted in progressive body weight loss. However, mice treated with combination of radiation and MXC-017 began to recover body weight starting day 3 post-irradiation **(Figure 4B)**. In contrast to DSS-induced colitis, radiation-induced enteropathy is characterized by distinct pathological features. Irradiated mice do not develop rectal bleeding, and colon length remains unchanged among the irradiated animals **(Figure 4C)**. As expected, radiation led to a breakdown of intestinal barrier integrity, as evidenced by significantly increased serum FITC-dextran levels. Again, addition of MXC-017 significantly attenuated this increase, indicating preservation of barrier function following irradiation **(Figure 4D)**. H&E staining of both the small intestine and colon revealed substantial radiation-induced epithelial injury, including disruption of normal tissue architecture, whereas MXC-017 treatment markedly reduced these pathological changes and preserved epithelial integrity **(Figure 4E)**. In addition, histological analysis further demonstrated that radiation induced significant shortening of small intestinal villi, which was partially mitigated by MXC-017 treatment **(Figure 4F)**. As in DSS-induced colitis, MXC-017 treatment also significantly reduced the radiation-induced expression of vimentin in colon tissues (**Supplementary Figure 2**).

The observed protective effects of MXC-017 against radiation enteropathy raised the question of whether MXC-017 could also protect cancer cells from radiation. To address this, we examined the effects of MXC-017 on the self-renewal capacity of prostate cancer stem cells using *in vitro* sphere-forming assays and extreme limiting dilution analysis (ELDA). Prostate cancer is believed to be organized hierarchically with a small population of cancer stem cells (CSCs) responsible for tumor initiation and maintenance. While several markers for exist that enrich for prostate cancer stem cells [32] we used functional assays of self-renewal to estimate their frequencies.

Castration-resistant human PC-3 and DU-145 prostate cancer cells were confirmed to express vimentin (**Figure 5A**). Treatment with MXC-017 alone significantly inhibited the sphere-forming capacity of PC-3 and DU-145 cells (**Figure 5B**). This effect was also observed in combination with radiation (**Figure 5B**) but did not reach statistical significance. An ELDA revealed that MXC-017 treatment significantly reduced the frequency of prostate cancer stem cells in PC-3 and DU-145 cells alone or in combination with radiation (**Figure 5C/D**). Lastly, MXC-017 treatment significantly reduced the migration of prostate cancer cells (**Figure 5E/F**) suggesting that MXC-017 may also have anti-metastatic effects in prostate cancer.

**Figure 5.**
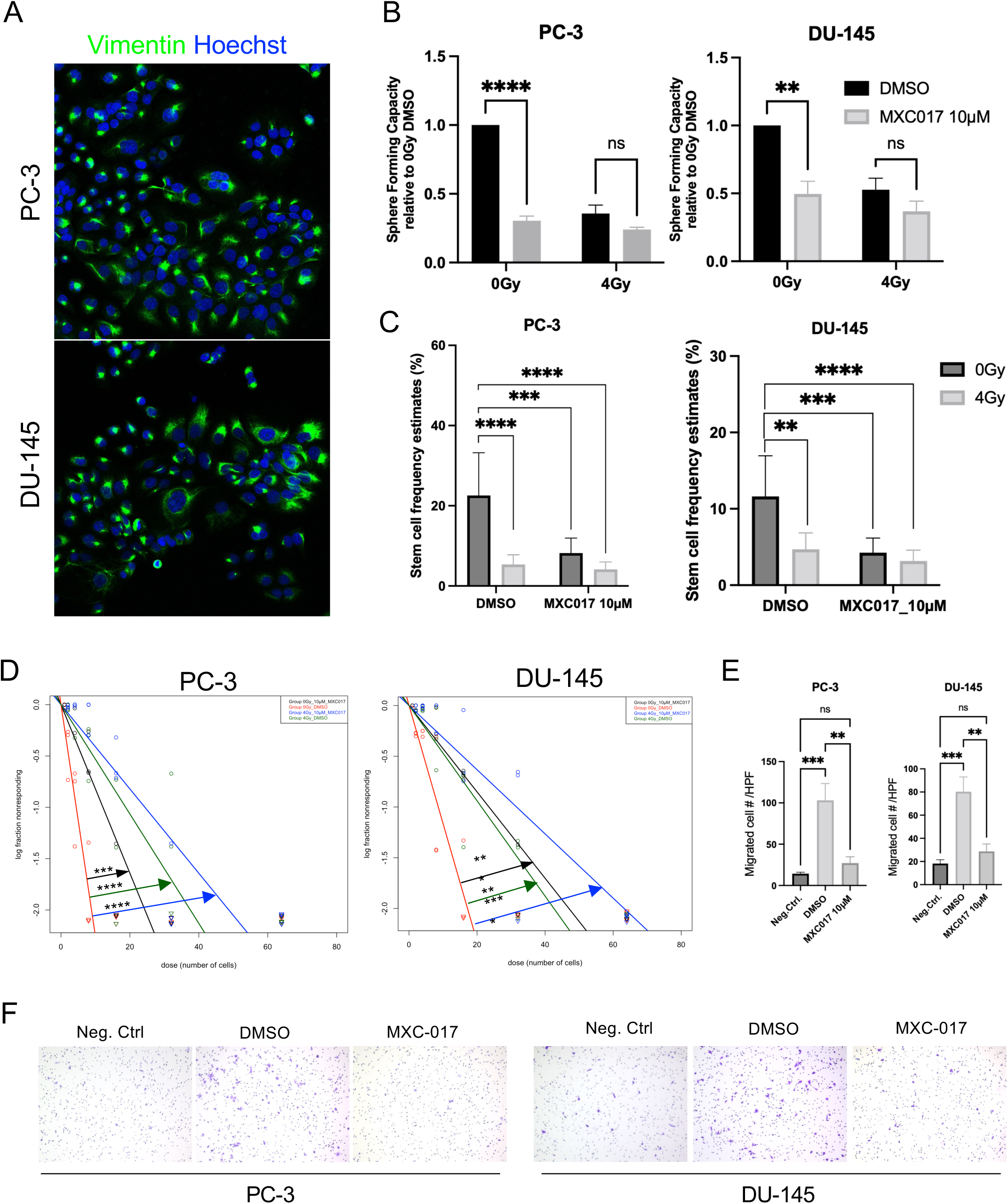
(**A**) Vimentin expression in human PC-3 and DU-145 prostate cancer cells. (**B**) Treatment with MXC-017 significantly inhibited the sphere-forming capacity of PC-3 and DU-145 cells. This trend was also observed in combination with radiation. (**C/D**) In extreme limiting dilution analyses MXC-017 treatment significantly reduced the frequency of prostate cancer stem cells in PC-3 and DU-145 cells alone or in combination with radiation. (**E/F**) MXC-017 treatment significantly reduced the migration of prostate cancer cells. 3-5 biological replicates were included in each assay. P-values were calculated using student t-test for B and C, One-way ANOVA for E. ** p-value < 0.01, *** p-value < 0.001, **** p-value < 0.0001.

## Discussion

UC is a chronic inflammatory condition of the colon and rectum characterized by periods of remission and exacerbation. Current treatments can control UC to some extent, but they do not cure it. Systemic treatments can cause substantial side effects and are not always effective. Therefore, novel therapeutic strategies for UC are needed.

Compared with the general population, patients with UC have a 2.4-fold higher relative risk of developing colorectal cancer (CRC) [33]. The standard of care for rectal cancer includes neoadjuvant chemoradiation followed by surgery. However, despite modern radiotherapy techniques this treatment is not well tolerated by patients with active UC. Moreover, radiotherapy is contraindicated in patients with severe UC, because radiation (but also chemotherapy protocols like FOLFOX) can exacerbate the disease. Similar restrictions apply to patients with other pelvic malignancies, such as prostate cancer or bladder cancer, and gynecological malignancies [34, 35].

MXC-017, originally developed to prevent radiation-induced cancer cell plasticity, is a novel compound that shows promise as a treatment for UC in the present study. MXC-017 was well tolerated in dose escalation studies in mice, and its maximum tolerated dose (MTD) was determined to be 150 mg/kg, with no signs of toxicity in the kidneys, liver, lungs. Likewise, MXC-017 administered at 150 mg/kg did not affect hematological or biochemical parameters [29].

Vimentin, the target of MXC-017 [29], is not primarily involved in the pathogenesis of UC. Rather, increased vimentin expression appears to be a consequence of inflammation and the associated tissue repair and remodeling, activation of the epithelial-to-mesenchymal transition and potential development of fibrosis. Vimentin knockout (KO) mice develop normally and exhibit a surprisingly subtle phenotype [36]. Consistent with this finding, MXC-017 treatment was well tolerated in wild-type mice with, despite locking vimentin intermediate filaments their assembled state [29].

Using a well-established mouse model of DSS-induced colitis, we observed symptoms typical of UC, including weight loss, rectal contraction, rectal bleeding, and diarrhea. MXC-017 treatment reduced the severity of colitis, preserved intestinal barrier function, and improved recovery. These findings are consistent with a previous study demonstrating reduced susceptibility to DSS-induced colitis in vimentin KO mice [37]. More recently, several other vimentin-binding or vimentin-stabilizing compounds have also been shown to be protect against DSS-induced colitis [38] while exhibiting anti-tumor activity [35, 39]. MXC-017 therefore expands this emerging class of vimentin-targeting compounds with potential therapeutic activity in both inflammatory disease and cancer.

Radiation enteropathy differs mechanistically from UC in that it is initiated not by disruption of the intestinal mucosal barrier but by a direct injury to the stem cell compartment of the intestinal epithelium. The subsequent loss of differentiated epithelial cells disrupts of the intestinal barrier function. In the small intestine, epithelial recovery begins approximately five days after irradiation, when regenerative crypts arise from surviving and reserve stem cells [40, 41]. MXC-017 does not have the properties of a classical radioprotector. Therefore, it is unlikely to act primarily by protecting the stem cell compartment of the intestinal epithelium from radiation. However, MXC-017 treatment prevented disruption of the intestinal epithelial barrier and preserved epithelial architecture, suggesting that it may support intestinal tissue integrity and recovery following irradiation.

These findings suggest that MXC-017 may have the potential to mitigate radiation-induced intestinal toxicity during radiotherapy for abdominal and pelvic malignancies. MXC-017 was originally developed as a treatment for glioma stem cells [29]. Notably, MXC-017 did not protect prostate cancer stem cells from radiation but instead enhanced the reduction in their frequency following irradiation. Together, these findings suggest further investigation of MXC-017 as an adjuvant to radiation therapy, with the potential to protect normal intestinal tissue without compromising, and potentially enhancing, antitumor efficacy.

The observed protective effects of MXC-017 on the intestinal epithelium and its antitumor activity are counterintuitive at first glance. However, under homeostatic conditions, vimentin expression and cytoskeletal dynamics are relatively limited in many differentiated epithelial cells, whereas tissue injury, inflammation, and malignant transformation can induce vimentin expression and remodeling. Pathological conditions such as IBD, radiation enteropathy, and cancer are associated with altered vimentin expression and filament dynamics. MXC-017 may therefore exert context-dependent effects by modulating the dynamic assembly and disassembly of vimentin filaments under conditions of cellular stress.

Our study has several limitations. Although MXC-017 treatment reduced the severity of radiation enteropathy in the small intestine of mice, we did not employ a mouse model of Crohn’s disease and therefore cannot draw conclusions about the compound’s efficacy in this form of inflammatory bowel disease (IBD). Furthermore, we did not investigate the long-term effects of MXC-017 treatment on the intestinal epithelium or determine whether MXC-017 protects the epithelium against subsequent DSS challenge. Finally, future studies should determine whether MXC-017, when combined with radiation, is effective in orthotopic mouse models of prostate cancer while also protecting the intestinal epithelium in mice with and without IBD.

We conclude that targeting vimentin with MXC-017 is a promising approach for treating UC and protecting both the colon and small intestine from radiation-induced toxicity. The observed antitumor activity of MXC-017 against prostate cancer stem cells is a novel finding that warrants further investigation.

## Author contributions

FP conceived of the study. XC, HD and MJ synthesized the MXC compound. LH and LA performed most of the experiments. CC, SJ, TP, and RS assisted in performing some experiments. FP, LH and LA analyzed the data. LH and FP drafted the manuscript. All authors contributed to and approved the final version of the manuscript.

## Data and Material Availability

All data are included in the article and/or *SI Appendix*.

---

**Supplementary Table 1.** Disease Activity Index (DAI) Scoring Criteria.

| **Parameter** | **Score** | **Criteria** |
|---|---|---|
| **Body weight loss (%)** | 0 | None |
|  | 1 | 1-5 % |
|  | 2 | >5-10 % |
|  | 3 | >10-15 % |
|  | 4 | >15% |
| **Stool consistency** | 0 | Normal |
|  | 2 | Loose stool |
|  | 4 | Diarrhea |
| **Rectal bleeding** | 0 | None |
|  | 2 | Mild bleeding |
|  | 4 | Gross bleeding |

**Supplemental Table 2.**
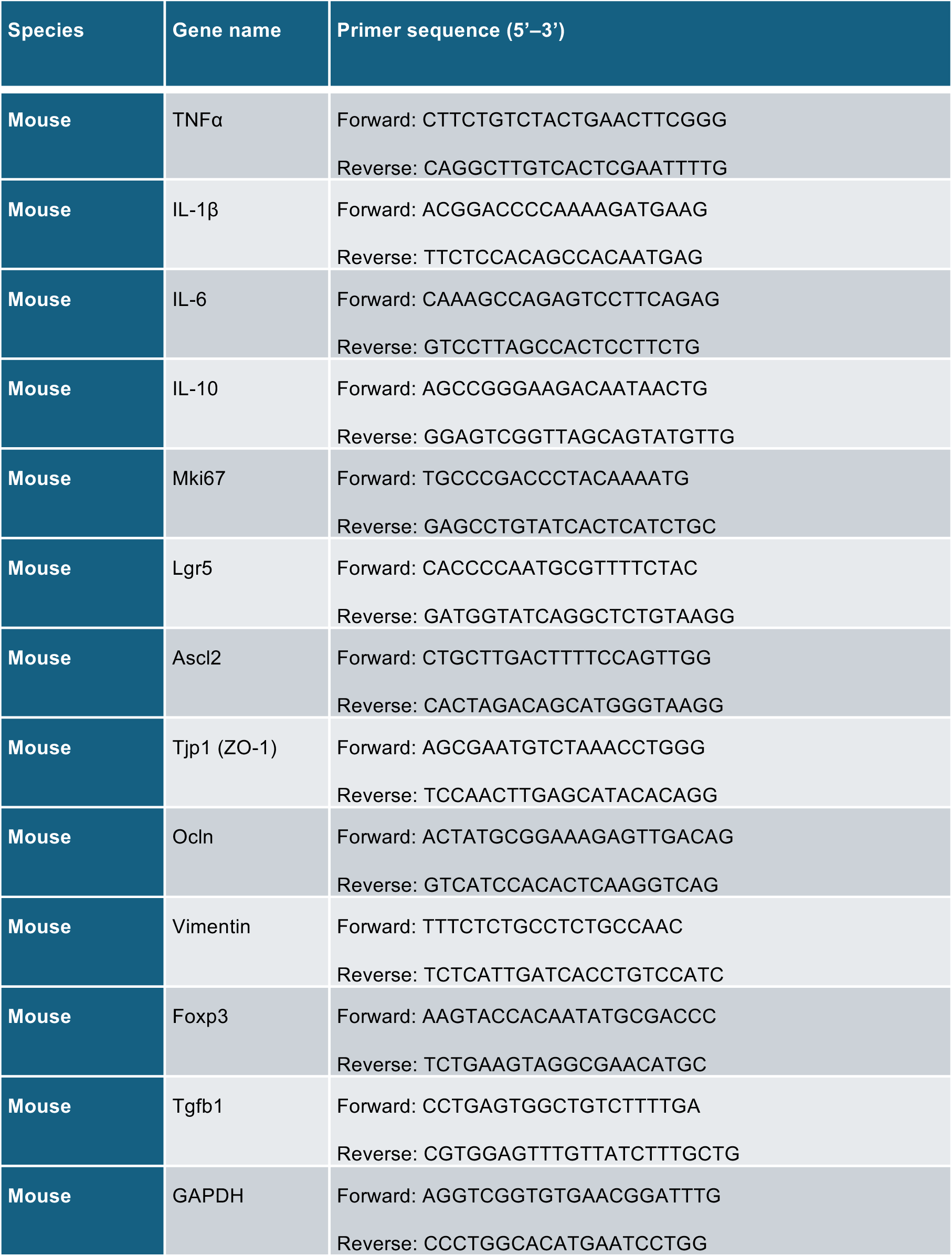
PCR primer sequences (Integrated DNA Technologies) used for RT-PCR experiments.

**Supplemental Figure 1.**
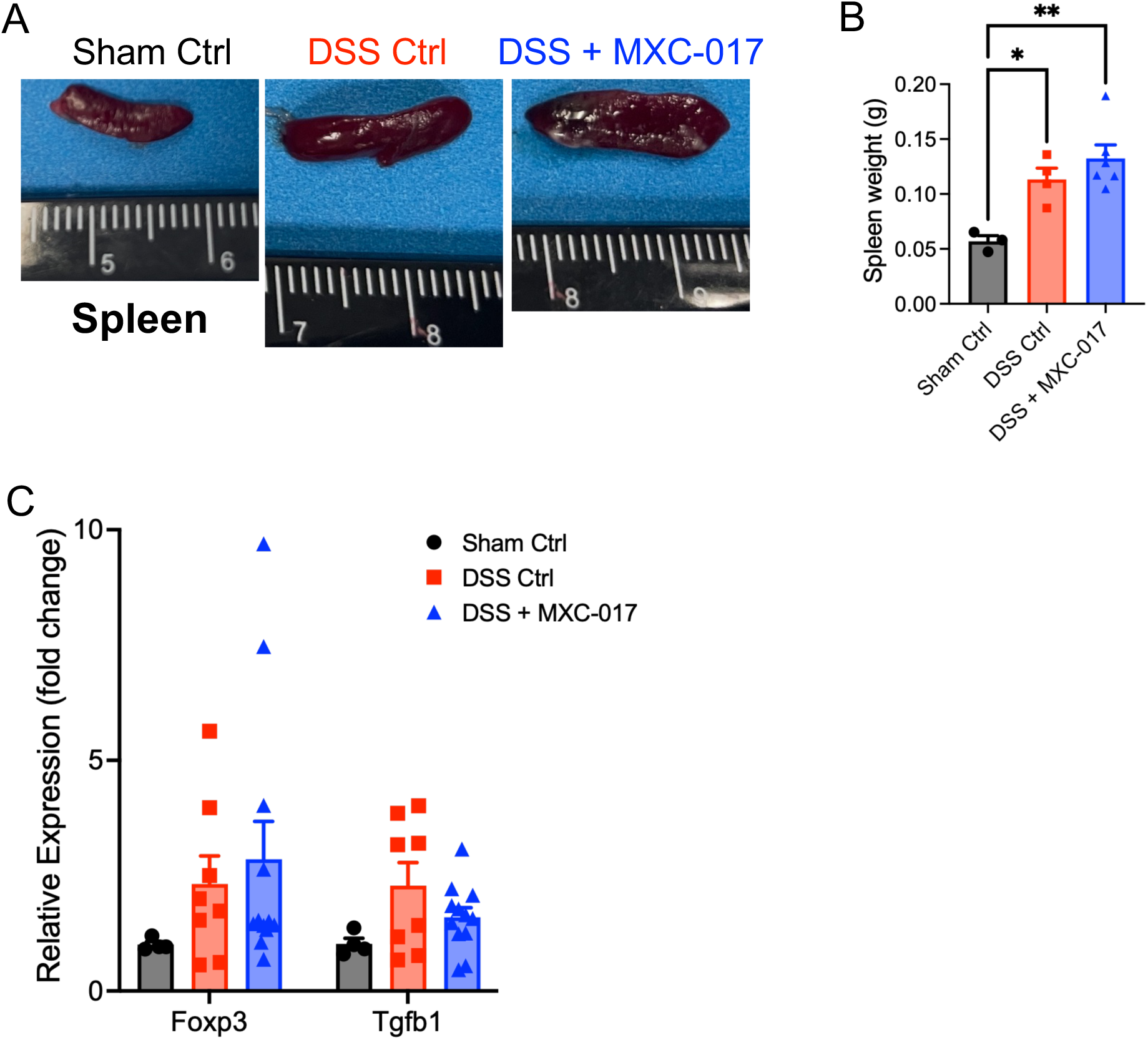

**Supplemental Figure 2.**
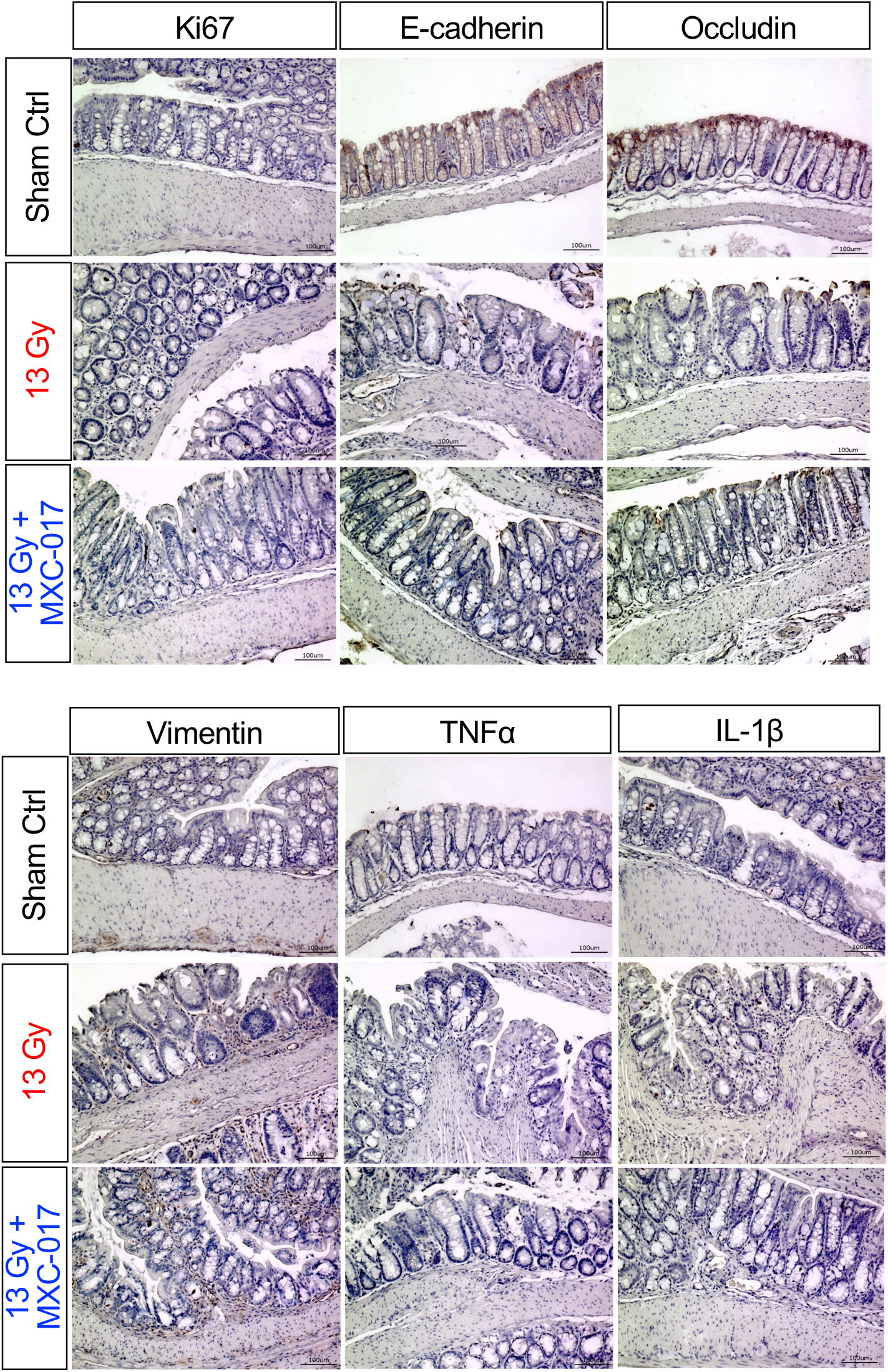

